# A Deep Dive into DNA Base Pairing Interactions Under Water

**DOI:** 10.1101/2020.06.04.134981

**Authors:** Rongpeng Li, Chi H. Mak

## Abstract

Base pairing plays a pivotal role in DNA functions and replication fidelity. But while the complementarity between Watson-Crick matched bases is generally believed to arise from the different number of hydrogen bonds in G|C pairs versus A|T, the energetics of these interactions are heavily renormalized by the aqueous solvent. Employing large-scale Monte Carlo simulations, we have extracted the solvent contribution to the free energy for canonical and some noncanonical and stacked base pairs. For all of them, the solvent’s contribution to the base pairing free energy is exclusively destabilizing. While the direct hydrogen bonding interactions in the G|C pair is much stronger than A|T, the thermodynamic resistance produced by the solvent also pushes back much stronger against G|C compared to A|T, generating an only ~1 kcal/mol free energy difference between them. We have profiled the density of water molecules in the solvent adjacent to the bases and observed a “freezing” behavior where waters are recruited into the gap between the bases to compensate for the unsatisfied hydrogen bonds between them. A very small number of water molecules that are associated with the Watson-Crick donor/acceptor atoms turn out to be responsible for the majority of solvent’s thermodynamic resistance to base pairing. The absence or presence of these near-field waters can be used to enhance fidelity during DNA replication.

**TOC Graphic:** 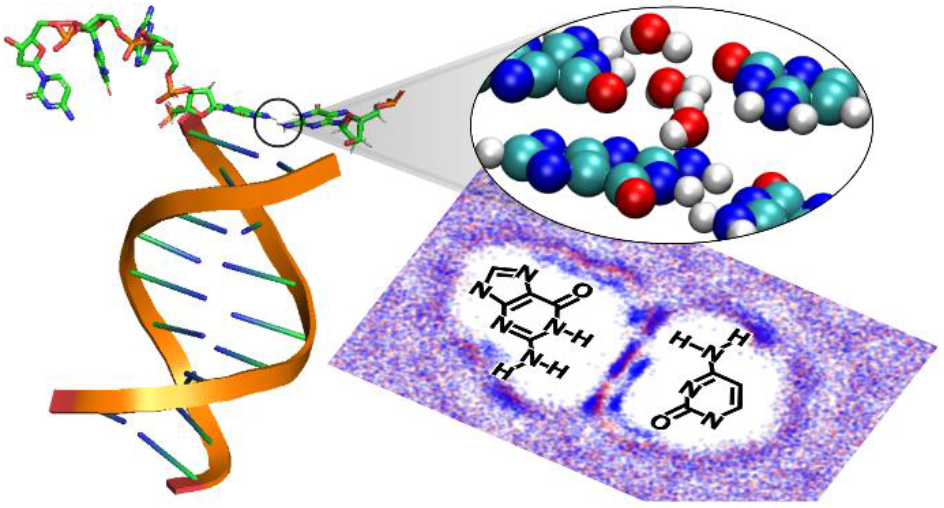

## 1. Introduction

DNA base pair complementarity is at the core of life. In Watson and Crick’s model of the double-helix DNA^1–3^, the molecular basis for precise replication and transmission of genetic instructions in the cell is founded on the complementarity between guanine (G) and cytosine (C) or adenine (A) and thymine (T) bases. In *The Double Helix*^4^, Watson described how the recognition that paired G|C and A|T bases in their dominant tautomeric forms (A, C amino, G, T keto) could form complementary hydrogen bonds with each other led to the final decisive clue to solving the puzzle of the structure of DNA, in a way that is consistent with Chargaffs rule^5^ and Franklin and Gosling’s X-ray diffraction profiles of B-DNA^6^.

Since Watson and Crick’s double helix model, hydrogen bonds have been invoked extensively in describing DNA physiology, both in the research literature as well as in many textbooks. For example, the common view that G|C base pairs, because they have three hydrogen bonds, are more stable that A|T pairs, which have only two, has become the basis for the standard IUPAC shorthand^7^ for G or C bases which are designated as “strong” (S), versus A or T bases which are designated as “weak” (W). While GC-rich regions on the genome are often found to be thermodynamically more stable than AT-rich regions, DNA melting studies^8^ suggest the difference may not be due to stronger hydrogen bonding between G and C – the melting free energy Δ*G* of a G|C pair stacked on top of another G|C pair being 3.1 kcal/mol, whereas it is 1.6 kcal/mol for a G|C stacked on top of A|T, compared to 1.9 kcal/mole for A|T on top of A|T. Thermodynamic studies at physiological temperatures^9^ also suggest that G|C pairs in DNA are only ~ −0.5 kcal/mol stable compared to A|T pairs, which are ~ +0.1 kcal/mol unstable, and the thermodynamic stabilities of DNA helices appear to be dominated instead by stacking free energies. These data seem to challenge the canonical view of the role of hydrogen bonding in DNA base pair complementarity^10^. Experimental results^11^ on shape analogs also suggest that bases that cannot hydrogen-bond can be replicated *in vivo* with high fidelity and other interactions may be key^12,13^.

What the original DNA base pairing model had not accounted for was the aqueous environment, but DNA biology operates in the context of water. While nucleobases can hydrogen bond with each other, water molecules can also form hydrogen bonds among themselves as well as with the nucleobases. The energy gained via the hydrogen bonds between DNA bases are partially offset by the loss of hydrogen bonding between them and nearby waters^14–16^. Recent spectroscopic studies of water clusters^17^ suggest that hydrogen bonds among waters are worth ~ 2 to 3 kcal/mol of free energy each. The loss of hydrogen bonding among water molecules could therefore compete against base pairing. Furthermore, solvated bases disrupt the solvent structure of water and can produce complex solvent-induced entropic effects^18–23^. Simulation studies have been performed to ascertain how hydrophobic solutes interact with the aqueous solvent, and Sharp, et al.^24,25^ and Duboué-Dijon et al.^26^ showed that hydrogen bond angle distribution varies the most while leaving the radial distribution of water almost unchanged. On the other hand, larger apolar solutes^27^ disrupt the hydrogen bonding network in water. Entropic forces arising from the aqueous solvent can therefore act against energetic factors produced by base pairing hydrogen bonds. Geometric fit between the paired bases can also take on extra importance under water due to the granularity inside the liquid structure of the aqueous environment^14–16^. Qi, et al^28^. reported the existence of two metastable states mediated by linking water molecules in a pairing pathway. It is proposed that different pairing geometries and separation distances can either include or exclude water molecules from the interface between two bases, causing a solvent-imprinted modulation of the hydrogen bonding interactions between them. Agris, et al.^29^ reported bridging water molecules for both canonical and modified bases in a molecular dynamics simulation. Lavery et al.^30,31^ have also used simulations to determine the pairing free energies of Watson-Crick pairs, and Chakraborty et al.^32^ have tried to determine the proper order parameters for a quantitative understanding of base pairing. Quantum chemistry calculations have also been performed in the presence of solvent waters to examine base complementarity^33^, and the possible role of secondary interactions between bases has also been suggested^34^.

In this paper, we focus on quantifying the intrinsic effects of water on free energy of a Watson-Crick base pair, with the goal of developing a theory to integrate this information into our solvent-free nucleic acid folding code^35^. Using large-scale Monte Carlo simulations of G|C or A|T pairs in water, we computed their pairing free energies as a function of separation. These free energy profiles provide clear evidence for the significance of the granularity of the liquid structure of the aqueous surroundings on base pairing. And since we are able to uniquely isolate the solvent’s contribution, we observe that while the direct interaction between G and C is much stronger compared to the A|T pair, the free energies of the G|C and the A|T pairs turn out to be similar under water, and the aqueous environment exerts a counteracting force that largely nullifies the energetic difference between the three hydrogen bonds in G|C and two in A|T. An analysis of the density profiles of solvent waters reveals a “freeze-thaw” behavior in the solvent as canonical Watson-Crick base pairs are formed.

## 2. Methods

Large-scale Monte Carlo (MC) simulations were carried out using rigid nucleobases with ideal geometries, solvated in a periodic box of approximately 4,000 TIP3P water molecules. The system’s total potential energy *U* is a sum of three terms:

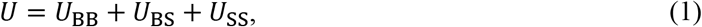

where they represent the base-base, base-solvent and solvent-solvent interactions. Amber ff99^36^ potentials were used, and electrostatic interactions were treated with an 8^th^ order fast multipole method (FMM) with Ewald summation^37^. The size of the simulation box was chosen large enough such that the bases had negligible interactions with any of their images. Each sample was equilibrated under constant pressure (1 atm) and constant temperature (310K, 320K, 345K). The use of MC permits straightforward umbrella sampling using hard walls, eliminating the restriction inherent in molecular dynamics simulations requiring strictly differentiable umbrella potentials.

Since standalone bases (i.e. with no sugar-phosphate backbone) were used in the simulations, the bases (A, G, C or T) were derived from the Drew-Dickerson dodecamer^38^ by cleavage at their respective N-glycosidic bonds. To avoid creating extra hydrogen bond donors at the cleavage sites, H9 atoms on A and G and H1 atoms on C and T were assigned zero partial charges, with excess charge on each base then re-assigned to the N9 atom for A and G or the N1 atom for C and T in order to maintain overall charge neutrality. The backbone was then discarded, and the bases were immersed in water and subjected to extensive equilibration. The standard Metropolis algorithm^39^ was used for random translations and rotations of the water molecules. For the free energy calculations, motions of the nucleobases were confined to either the stretch, the buckle or the propeller direction (see Fig. 1). Umbrella sampling^40^ was employed to compute the free energy and to ensure adequate sampling across the entire free energy profile. Umbrella windows in the stretch direction were each 0.4 Å wide, and they span interbase separations between 2.6 to 8 Å, measuring by the distance between hydrogen bonding donor and acceptor between bases. In order to reliably align free energies across all windows, nearest-neighbor umbrella windows had 0.2 Å overlaps with each other, equal to half the width of each window. Additionally, the forces acting on the bases from the gradients of *U*_BB_, *U*_BS_ and *U*_SS_ in the stretch direction was also evaluated during the simulations to facilitate an independent calculation of the reversible work.

**FIGURE 1.**
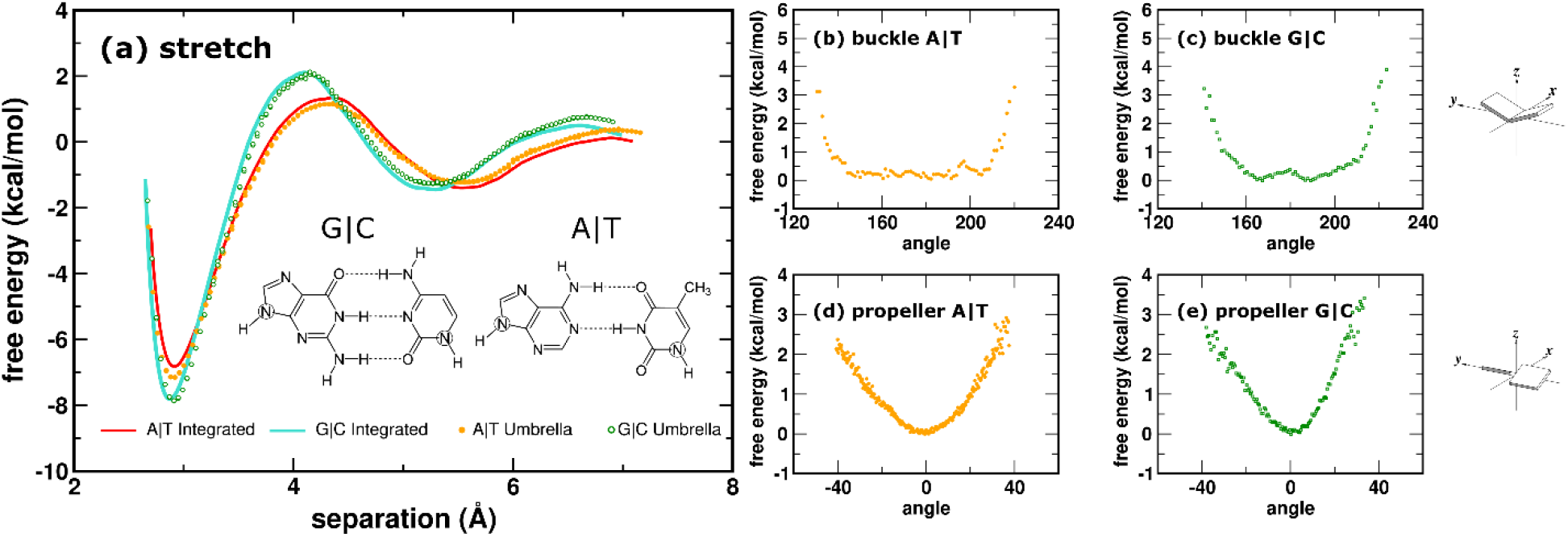
(a) Free energy profiles Δ*G*(*x*) for the A|T and G|C pairs as a function of interbase separation *x* in the stretch direction. Solid red line and closed orange circles show A|T free energy calculated using umbrella sampling and the integral of the reversible work, respectively. Solid cyan line and open green circles show the same for G|C. Geometries of the base pairs are shown in the legend, where the stretch direction is parallel to the dotted lines, and glycosidic N atoms are circled. (b) Free energy for A|T in the buckle direction orthogonal to the stretch. (c) Free energy for G|C in the buckle direction. Cartoon on the top right illustrates buckle motion. (d) and (e) Free energies for A|T and G|C in the propeller direction. Cartoon on lower right illustrates propeller motion. Open circles in each panel show typical statistical uncertainties in the free energy data.

The computational model has been designed to quantify the role of water in the aqueous environment on the free energy of a Watson-Crick base pair. These calculations specifically do not include the sugar-phosphate backbone or any protein. The goal is to ascertain how much of the complementarity interactions between nucleobases could be attributed just to water, and how many and which water molecules in the vicinity of the bases are the primary effectors. Nucleobases in this calculation have been treated as rigidly planar. While planarity of the nucleobases has generally been assumed, there are reports^41^ questioning their rigidity specifically at glycosidic N atom positions (N1 in pyrimidines, N9 in purines), but simulations focusing on base pairing effects^30^ have assumed chemical bonds on the bases can be constrained as rigid. Counterions are important to the stability of nucleic acids due to the high density of negative charges on the sugar-phosphate backbone^42–44^, but the present model does not include the backbone or any added salt.

## 3. Results

### G|C and A|T pairs have similar free energies under water

The calculated free energies of G|C and A|T base pairs in water are shown in Fig. 1. Fig. 1(a) shows free energies Δ*G*(*x*) as a function of separation *x* between the bases along the stretch direction. For each pair, Δ*G*(*x*) was calculated two different ways to assess the accuracy of the computed values. The G|C free energy computed directly from umbrella sampling are shown in Fig. 1(a) as the open green circles, and for A|T as the closed orange circles. The integral of the reversible work, which is the ensemble-average gradient of the system’s potential energy *U*, of bringing two infinitely-separated bases together provides a second independent method for calculating the free energy. This is shown in Fig. 1(a) as the cyan line for G|C and the red line for A|T. The agreement in Fig. 1(a) between these two methods indicates that the accuracies of the results are better than ~ 0.2 kcal/mol for both G|C and A|T. At the minimum free energy separation of *x* ~ 2.8 Å, the G|C pair is approximately −8.0 kcal/mol stable relative to unpaired G and C, whereas the A|T pair is approximately −6.9 kcal/mol stable. Noticeably, the stabilities of the G|C and A|T pairs are quite similar, and they do not appear to directly correlate with the expected number of hydrogen bonds between the bases (three in G|C, two in A|T). Each umbrella calculation in Fig. 1(a) computes the free energy change as a function of bringing a base in reversibly from infinite separation into pairing geometry with another base. Since the free energy *G* is a thermodynamic function, its change Δ*G* is path-independent; therefore, the calculated Δ*G* between paired (at the minimum of the free energy profiles) versus unpaired bases (at infinite separations) is not affected by any constraint applied in the stretch direction.

Upon closer examination of the free energy profiles Δ*G*(*x*) for both G|C and A|T, Fig. 1(a) reveals an unexpected nonmonotonic dependence on the separation *x* between the bases if the attraction between the bases was due to direct hydrogen bonding interactions. The global minimum free energy separations in G|C and A|T pair are *x* ~ 2.8 and 2.9 Å, respectively. But a second local minimum in Δ*G*(*x*) is observed at an interbase separation of *x* ~ 5.3 Å for G|C and 5.5 for A|T. Between the global and second minima, a maximum is located approximately midway between them in each pair. Oscillatory variations in Δ*G*(*x*) are *not* expected if the two bases within each pair interact purely via direct hydrogen bonding interactions. These nonmonotonic variations in Δ*G*(*x*), as will be shown below, turn out to be signatures of the underlying granularity of the liquid structure of the aqueous solvent. The global minimum and the second minimum in the profiles of both G|C and A|T are separated by ~ 3 Å, coinciding roughly with the diameter of one water molecule. Similar G|C and A|T pair free energy profiles have also been reported previously by Lavery et al.^45,30,31^ but they included rotations of the bases, whereas in our results in Fig. 1(a), the two bases have been constrained to stay coplanar.

To assess the effects of rotations separately, we then computed the free energy profiles for these motions orthogonal to the stretch direction at the global minimum free energy distance. The buckle free energies for A|T and G|C are shown in Figs. 1(b) and (c). The propeller free energies are shown in Figs. 1(d) and (e). Clearly, the curvatures of these free energy profiles in both the buckle and propeller directions exert strong restoring forces for each base pair to return to its preferred coplanar geometry. These free energy profiles were calculated by monitoring the frequencies of occurrence of different angles between the two bases in the ensemble.

### Aqueous environment counteracts stronger direct interaction in G|C than A|T

In addition to Watson-Crick pairs, we have computed the free energy of a G|T pair. The result is shown in Fig. 2(a) as the open blue diamonds, compared to A|T (closed orange circles) and G|C (open green squares) results from Fig. 1(a). The “idealized” approach geometry of the G|T pair employed for this calculation is shown in the legend. The two donor-hydrogen-acceptor (D-H-A) axes (the dashed lines between G and T in Fig. 2(a)) are approximately parallel to each other, similar to the two D-H-A axes in A|T (see Fig. 1(a)). As such, the thermodynamic stability of the G|T pair turns out to be very similar to the A|T pair, ~ −6.8 kcal/mol. The free energy profile Δ*G*(*x*) of G|T exhibits similar solvent-induced oscillations as A|T and G|C, though the variations are somewhat muted. Previous molecular dynamics results for A|U and modified A|U base pairs^29,46^ also showed similar robustness against modifications, which agree with our observations. However, it is important to stress that the sugar-phosphate backbone or DNA polymerase would probably have prevented the G|T pair to adopt this idealized approach geometry during DNA replication, and geometry, shape or fit could be key factors in deciding the energetics of a G|T mismatch pair versus canonical pairs.

**FIGURE 2.**
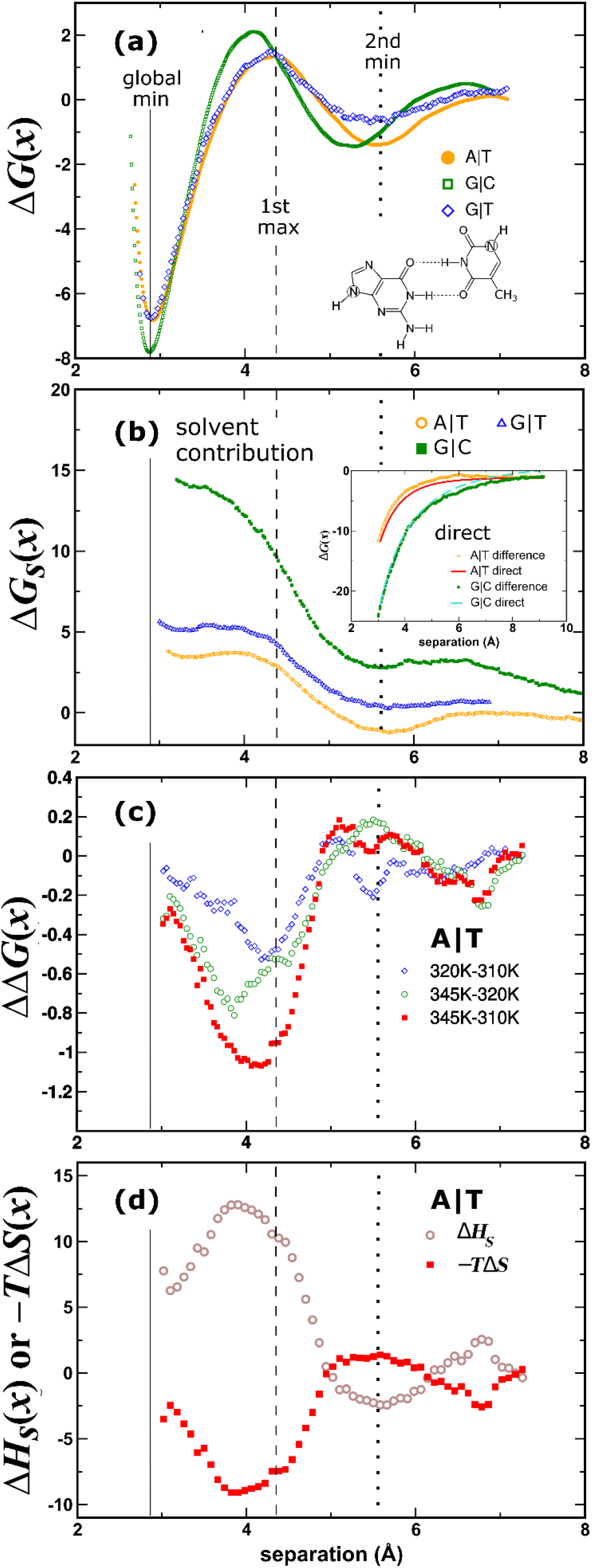
(a) Free energy profiles Δ*G*(*x*), in units of kcal/mol, for A|T (closed orange circles), G|C (open green squares) and the G|T wobble pair (open blue diamonds), compared against each other, with A|T and G|C data from Fig. 1(a). The geometry of the G|T pair is shown in the legend, with the dashed line defining the stretch direction and glycosidic N atoms are circled. The vertical dotted line indicates position of the second local minimum in the A|T free energy profile (orange circles), the vertical dashed line, the position of the first A|T maximum, and the solid line, the position of the global minimum. Open circles show typical statistical uncertainties in the umbrella free energies. (b) The solvent’s contribution, Δ*G*_s_(*x*), to the A|T (open orange circles), G|C (closed green squares) and G|T (open blue triangles) pairing free energies, as a function of interbase separation *x*. Inset: The difference between Δ*G*(*x*)−Δ*G*_S_(*x*) corresponds to the direct base-base interactions for A|T (open orange circles) and G|C (close green squares). These are compared against the direct interaction energies between A|T (solid red line) and G|C (solid cyan line). (c) Differences between free energy profiles, ΔΔ*G*(*x*), computed at three different temperatures, 310K, 320K and 345K, for the A|T pair. (d) The entropic component, −*T*Δ*S*(*x*), in closed red squares, and the enthalpic component, Δ*H_S_*(*x*), in open brown circles, of the solvent’s contribution Δ*G_S_*(*x*) to the pairing free energy of A|T. The vertical dashed line through all the panels marks the *x* position of the first maximum in the A|T pair free energy Δ*G*(*x*), whereas the dotted line marks the position of the second minimum. See Fig. 4 for the arrangement of water molecules in the vicinity of the bases.

To understand the role of the aqueous solvent in shaping the free energies of the base pairs, we extracted the solvent’s contribution to the free energy, using two independent methods. First, since in Eq. (1) the potential energy *U* of the entire system is decomposable into *U*_BB_ + *U*_BS_ + *U*_SS_, the solvent’s contribution to the force between the two bases is entirely captured by the gradient of 〈*U*_BS_ + *U*_SS_〉 in the stretch direction *x*, where the brackets indicate an ensemble average. The integral of 〈∇*U*_BS_ + ∇*U*_SS_〉 with respect to *x*, which is the reversible work of stretching the base pair, then yields the solvent’s contribution to the pairing free energy. Denoting this solvent contribution to the free energy by Δ*G*_S_(*x*), the free energy of the base pair becomes Δ*G*(*x*) = Δ*G*_S_(*x*) + Δ*G*_D_(*x*), where Δ*G*_D_(*x*) is the “direct” interaction between the two bases. A second way of computing Δ*G*_S_(*x*) is to perform umbrella sampling on a system with potential *U*_BS_ + *U*_SS_ only. To verify the accuracy of the computed Δ*G*_S_(*x*), we cross-checked these two sets of calculations against each other. Their numerical results agree to within 0.2 kcal/mol, giving us high confidence in the computed Δ*G*_S_(*x*) values.

Fig. 2(b) shows the solvent’s contribution, Δ*G*_S_(*x*), to the pairing free energy of the A|T pair in open orange circles, for the G|C pair in closed green squares and for the G|T pair in open blue triangles. Interestingly, for all three pairs, the results in Fig. 2(b) suggest that the solvent’s contribution to the pairing free energy is almost exclusively destabilizing over almost the entire range of interbase separations *x*. In the rest of the paper, we adopt a reference where two infinitely-separated bases are *defined* to have both Δ*G*(∞) = 0 and Δ*G*_S_(∞) = 0. This is because two well-separated bases should have no interaction with each other, whether in the direct energy between them or the free energy induced by the solvent. The A|T and G|C data in Fig. 2(b) extend to separations well beyond the right side of the graph, illustrating this expected asymptotic reference behavior for Δ*G*_S_(*x*) at large separations.

In Fig. 2(a) the *x* positions of the global minimum, the first maximum and the second minimum are marked by three vertical line (solid, dashed and dotted, respectively), and these markers are reproduced on all the other panels in Fig. 2. Fig. 2(b) shows that the solvent’s contribution to the A|T free energy (open orange circles) produces a ~ +4 kcal/mol destabilization of the A|T pair relative to the reference state. And as A and T come together to pair, the solvent counters their pairing by imposing a free energy penalty according to Δ*G*_S_(*x*), which is almost entirely repulsive. Comparing Δ*G*_S_(*x*) to the free energy Δ*G*(*x*) in Fig. 2(a) which shows that A|T is overall stable by −6.8 kcal/mol, this suggests that the naked direct hydrogen bonding interactions Δ*G*_D_(*x*) should have a raw strength of ~ −6.8 - (4) = −12.8 kcal/mol for A|T. To confirm this, the difference Δ*G*(*x*) – Δ*G*_S_(*x*) is calculated and shown in the inset in Fig. 2(b) as the open orange circles for the A|T pair, and as expected this difference is identical to the direct interaction Δ*G*_D_(*x*) between the bases in the absence of solvent, which is shown as the red solid line. The naked direct hydrogen bonding interaction between A and T exhibits a smooth monotonic increase in the attraction between them as they come together to pair. The solvent is therefore solely responsible for the oscillations observed in the A|T pairing free energy in Fig. 2(a).

Turning to the G|C pair, its results are largely consistent with A|T. But in the case of G|C, the solvent seems to push back much more forcefully against the direct interaction between G and C, compared to A|T. Fig. 2(b) shows Δ*G*_S_(*x*), the solvent’s contribution to the G|C pairing free energy, as the green squares. As G and C come together to pair, the solvent counters their pairing by imposing a free energy penalty between well separated G and C and the global minimum free energy of approximately +13 kcal/mol. In contrast, this was +4 kcal/mol for A|T. The naked direct hydrogen bonding interaction between G|C is shown in the inset of Fig. 2(b) as the closed green circles (Δ*G*(*x*) – Δ*G*_S_(*x*)) and the cyan line. The magnitude of the direct hydrogen bonding interactions in G|C is clearly much stronger than A|T since there are three hydrogen bonds in G|C but only two in A|T. However, the solvent also seems to produce a disproportionately stronger resistance against pairing in G|C than in A|T, so that the thermodynamic stability of G|C ends up being only ~ 1.2 kcal/mol more stable than A|T as Fig. 2(a) suggests.

The results in Fig. 2(b) demonstrate that the aqueous environment exerts a counteracting free energy penalty against the direct attractive interaction between the bases. This thermodynamic resistance offered by the solvent seems to be much stronger in G|C than A|T, acting to push back on the stronger direct hydrogen bonding interactions in the G|C pair. The net result of this is a G|C pairing free energy that is only marginally more stable than A|T, and the thermodynamic stabilities of G|C and A|T do not scale with the expected number of hydrogen bonds in them.

Why is the solvent’s free energy Δ*G*_S_(*x*) always destabilizing, and why is the strength of the solvent’s resistance so different between A|T and G|C? The results in Fig. 2(b) suggest that the solvent is most stable when the bases are unpaired but sacrifices free energy when the bases pair. While this can either be enthalpic or entropic in origin (or both), one can postulate that when bases are well-separated, the solvent waters may be better able to thermodynamically satisfy the hydrogen-bonding donor(s) and acceptor(s) on the two individual bases and among themselves. Fig. 2(b) suggests that when the bases pair, water molecules in their vicinity may no longer be able to maximally satisfy all the hydrogen bonds, either within the solvent or between the solvent and the bases. This “hydrogen-bonding deficit” that is generated in the aqueous solvent then leads to the free energy increase exhibited in Δ*G*S(*x*). Enthalpy and entropy data to be presented below confirm that the thermodynamic cost incurred by the solvent do indeed vary with the number of water molecules captured between the bases. Regarding the magnitude of these thermodynamic penalties, Fig. 2(b) shows that a G|C pair costs the solvent ~ +13 kcal/mol, whereas A|T and G|T are much milder, at ~ +4 kcal/mol and +5 kcal/mol. Based on these data, it appears that the free energy cost to the solvent is not proportional to the number of hydrogen-bonding donor and/or acceptor atoms that are exposed to water when the bases come apart. However, the A|T and G|T pairs, both having two direct hydrogen bonds, appear to cost the solvent approximately the same free energy penalty, whereas for G|C, which has three direct hydrogen bonds between them, costs the solvent substantially more. Enthalpy and entropy data below will show that there is a complex relationship between these thermodynamic functions and the number of water molecules that could be trapped between the bases.

### Comparison with experimental free energies and the interplay between stacking and pairing

Each of the pairing free energies presented in Figs. 1(a) or 2(a) as a function of the donor-acceptor atom separation is for a single pair of bases constrained to stay coplanar with each other when the distance between them is being stretched. Along this degree of freedom, we have seen that the pairing free energy is Δ*G* −8.0 kcal/mol for G|C and −6.8 kcal/mol for A|T. At first sight, these calculated values appear to disagree with the experimental measurements summarized in the Introduction. There are two reasons for this. First, the free energy profiles in Fig. 1(a) and 2(a) are for the stretching direction only, and there are additional components from the other orthogonal degrees of freedom (buckle and propeller). In the limit of infinitely separation, either unpaired base has no overall preferred orientation. If we consider two separated bases as the “product” state of an unpairing reaction, then in the “reactant” state where the two bases are paired, they are subjected to additional rotational free energy penalties as Figs. 1(b) through (e) show. In the reactant state, the buckle and propeller angles are confined to very limited ranges, but in the product they are unconstrained. We can estimate the additional free energy costs associated with these orthogonal degrees of freedom by calculating the thermally accessible range of buckle and propeller angles at 37°C. For the A|T pair, the rotational free energy profiles suggest an extra free energy cost of ~ +1.0 kcal/mol in the buckle and +1.3 kcal/mol in the propeller mode. Whereas for G|C, they are ~ +1.1 kcal/mol and +1.4 kcal/mol, respectively. Accounting for these, the additional free energy penalties incurred in bringing the bases into a coplanar geometry in order for them to pair requires another ~ +2.5 kcal/mol for A|T and ~ +2.7 kcal/mol for G|C under water.

There is a second reason why the experimentally measured free energies do not correspond directly to those in Fig. 1(a) and 2(a). Because DNA is a ribose-phosphate polymer, there is a reduction in the conformational entropy of the polymer backbone when bases come together to pair. This conformational entropy has been estimated to incur an additional free energy cost ~ +5.1 kcal/mol when a new base pair is formed stacked on top of another base pair^47^. Accounting for these two additional sources of free energy costs, the apparently stable −6.8 kcal/mol A|T pairing free energy in Fig. 1(a) in the stretch direction becomes only +0.6 kcal/mol, which renders it slightly unstable. On the other hand, the G|C pairing free energy, while having a free energy in the stretch direction of −8.0 kcal/mol, becomes only −0.4 kcal/mol stable when all other free energy factors are taken into account. These estimates are very consistent with the experimental measurements of Yakovehuk et al.^9^, which suggest that the G|C base pair is only marginally stable thermodynamically, while the A|T pair is slightly unstable.

To completely conform with experimental conditions, one final factor must be included. Inside a DNA duplex, it is essentially impossible to separate the effects of base pairing from stacking, since one base pair is always stacked on top of another inside B-form DNA. Additionally, from the point of view of DNA replication, copy-strand DNA is synthesized by polymerase extending the primer one nucleotide at a time in the 5’ to 3’ direction, by selecting the correct dNTP complementary to the template strand to form a new base pair stacked on top of the previous one. Under these conditions, the most relevant pairing free energy is not the quantity between one A and one T or between one G and one C, but instead, should be the free energy for adding one new base to make the copy *with its template base already stacked on top of another base pair.* For such base pair “doublets”, there are ten distinct combinations, e.g. pairing G to a C on top of G|C (denoted GG|CC), pairing G to a C on top of C|G (CG|GC), pairing G to a C on top of A|T (AG|TC), etc. For the “singlet” pair, there are only two combinations (A|T, G|C) and their free energies have been considered above. To completely understand base pairing under water, it is necessary to also consider the free energy of formation of the doublets. The computational work involved in calculating the free energies of all doublet combinations is expensive. Instead, we have selected one example to quantify the effects of stacking.

In Fig. 3(a), we show the pairing free energy profile for the GG|CC doublet, for adding a C to a G on top of an already formed G|C pair, as the solid purple line. As in the singlet calculations, the G being added was constrained to be coplanar with the G. The solid purple line in Fig. 3(a) shows the GG|CC pairing free energy Δ*G*(*x*) as a function of separation *x*, compared to the same result for the G|C singlet from Fig. 2(a) as the dashed green line. Fig. 3(a) shows that the global minimum free energy donor-acceptor distance in the GG|CC doublet remains ~ 2.8 Å, similar to singlet G|C. For the GG|CC doublet, the pairing free energy is deeper, at ~ −13.8 kcal/mol, compared to −8.0 kcal/mol for G|C. Because stacking provides additional stabilization for the newly added G, the stronger Δ*G* for GG|CC is expected. Fig. 3(a) suggests that GG|CC is ~ −5.8 kcal/mol more stable than singlet G|C, which is comparable to the previously reported stacking free energy of G upon G^20^.

**FIGURE 3.**
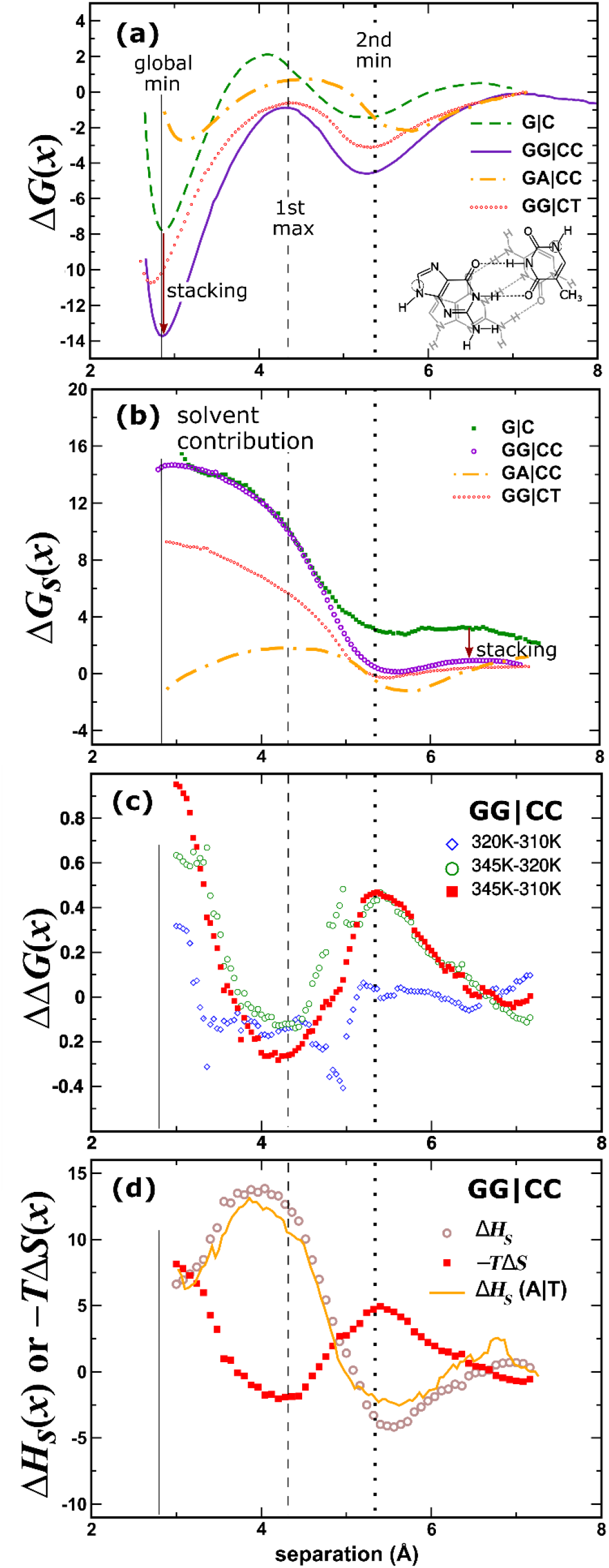
(a) Free energy profiles Δ*G*(*x*), in units of kcal/mol, for GG|CC (solid purple line), GG|CT (red dotted line) and GA|CC (dotted dashed orange line), compared to G|C (dashed green line). The vertical dotted line indicates position of the second local minimum in the GG|CC free energy profile, the vertical dashed line, position of the first GG|CC maximum and the vertical solid line, the global minimum. The red down arrow indicates the stacking free energy contribution to the GC pair upon another GC. Legend shows geometry of the G|T pair (black) below the G|C pair (gray) in the GG|CT doublet. Open circles show typical statistical uncertainties in the umbrella free energies. (b) The solvent’s contribution, Δ*G_S_*(*x*), to the GG|CC (open purple circles), GG|CT (red dotted line), GA|CC (dotted dashed orange line) and G|C (closed green squares) free energies, as a function of interbase separation *x*. The difference between G|C and GG|CC is due to the stacking free energy, indicated by the red arrow. (c) Differences between free energy profiles, ΔΔ*G*(*x*), computed at three different temperatures, 310K, 320K and 345K, for the GG|CC pair. (d) The entropic component, −*T*Δ*S*(*x*), in closed red squares, and the enthalpic component, Δ*H_S_*(*x*), in open brown circles, of the solvent’s contribution Δ*G_S_*(*x*) to the pairing free energy of GG|CC. The enthalpic component for A|T is shown as the orange line as a comparison. See Fig. 4 for the arrangement of water molecules in the vicinity of the bases.

Earlier, we have shown that the free energy of a G|C base pair becomes only marginally stable at ~ −0.4 kcal/mol after accounting for the free energy costs associated with the restricted rotations of the bases and the conformation entropic penalty for the sugar-phosphate backbone. Now adding stacking to this, experimental estimates suggest that the extra stability coming from stacking for a GC pair on top of another GC pair should be ~ −2.0 kcal/mol^9^. However, our stacking results seem to suggest that this is ~ −6.0 kcal/mol for GG|CC. To resolve this apparent discrepancy, notice in the case of doublet GG|CC that the buckle and propeller rotations of the G being added to form the new base pair are much more constrained compared to in singlet G|C, because the newly added G must lie flat against the base pair underneath to facilitate maximal stacking. However, rotational free energies in the doublet were difficult to evaluate numerically because the ranges of the accessible buckle and propeller angles were quite narrow. From the simulated configurations, we estimated that the accessible range of buckle angles was about 5 times narrower in GG|CC compared to G|C and the propeller angle was close to 20 times narrower. Based on these estimates, the rotational free energy costs to forming GG|CC is expected to be ~ +3 kcal/mol *higher* in GG|CC than G|C, and this reduces the apparent stacking free energy contribution to a GG|CC doublet from ~ −5.8 kcal/mol to only −2.8 kcal/mol, which puts it roughly in line with experimental measurements^9^.

The pairing free energy for the GG|CT doublet, with a mismatched G|T pair, is shown in Fig. 3(a) as the red dotted line. The legend shows the geometry of the G|C pair in gray above the G|T pair in black, inside this GG|CT doublet. The free energy shown is for adding a T, coming in coplanar, to pair with a G that is already stacked below a G|C. Compared to the singlet G|T pair in Fig. 2(a), Fig. 3(a) shows that the GG|CT doublet is more stable by ~ 4 kcal/mol. The free energy for the GA|CC doublet, with a mismatched A|C pair, is also shown in Fig. 3(a) as the dotted dashed orange line. A C brought in to pair with an A already stacked against a G|C pair turns out to be unstable. Most of the uphill free energy in this case is due to unfavorable electrostatic interactions between C and A, which can only make one hydrogen bond in the same pairing geometry as in G|T. This suggests that under conditions representative of replication, a A|C mismatch should be readily recognizable based on this free energy discrepancy. However, this may be sensitive to the specific choice of the force field employed because small differences in the partial charges assigned to the base atoms can alter the GA|CC results. But for the GG|CC doublet, the data are much more robust against force field variations because the free energy minimum there is quite deep. Also expected to be robust are the GG|CT results, which unlike an A|C mismatch on top of G|C, show that a G|T mismatch on top of G|C is stable. But compared against the matched G|C in GG|CC, a mismatched G|T in GG|CT is ~ +3 kcal/mol less stable, even though Fig. 2(a) shows that the G|T singlet is only ~ +1 kcal/mol less stable than G|C. This suggests that the exclusion of solvent molecules by adding more stacked base pairs could diminish the solvent’s resistance, and this effect appears to be disproportionately higher for mismatched base pairs.

Fig. 3(b) shows the solvent’s contribution Δ*G*_S_(*x*) to the pairing free energy of the GG|CC doublet in open purple circles. The same data for the G|C singlet from Fig. 2(b) are shown in closed green squares for comparison. Similar to all singlet pairs in Fig. 2(b), Fig. 3(b) shows that the solvent’s contribution is exclusively destabilizing for GG|CC. The thermodynamic resistance exerted by the solvent against base pairing in the GG|CC doublet is almost identical to the G|C singlet at ~ +13 kcal/mol. Assuming the effects of base pairing and base stacking are additive, the difference between Δ*G*_S_(*x*) for GG|CC (purple circles) and G|C (green squares) should correspond approximately to the solvent’s contribution to the stacking free energy, which is indicated by the downward arrow in Fig. 3(b), ~ −3 kcal/mol. Comparing this to the full free energy difference of −5.8 kcal/mol between GG|CC and G|C from Fig. 3(a), this suggests that the solvent is responsible for the vast majority of the thermodynamic force behind base stacking, consistent with recent simulation^20^, theory^35^ and experiment^48^.

The solvent’s contribution Δ*G*_S_(*x*) to the pairing free energy for the GG|CT doublet with a mismatched G|T is shown in Fig. 3(b) as the red dotted line. Compared to the GG|CC doublet with a matched G|C pair (open purple circles), the solvent’s contribution is weaker with the mismatched G|T by a significant amount of ~ 6 kcal/mol. The free energy reveals the same solvent-imprinted signature at the position indicated by the vertical dotted line. But comparing this to the GA|CC doublet with a mismatched A|C pair, shown as the orange dotted dashed line in Fig. 3(b), the solvent’s contribution is even weaker with the A|C mismatch. These GG|CC, GG|CT and GA|CC Δ*G*_S_ results suggest a possible compensation effect could exist between the strength of the direct interaction between the bases versus the solvent’s free energy. When the direct base-base interaction is strong, such as in the case of GG|CC, the solvent pushes back more forcefully, but when the direct base-base interaction is weaker, such as in the case of GG|CT or GA|CC, the solvent’s resistance is concomitantly milder.

In addition to 310K, the same free energy calculations had been performed at 320K and 345K for GG|CC. From those results, we present ΔΔ*G* data among the three temperatures for A|T in Fig. 2(c) and for GG|CC in 3(c). Using the mean finite difference ΔΔ*G*/Δ*T* measured for the three temperature gaps, we derive an estimate for the entropic component of Δ*G*, −*T*Δ*S*, for each pair in Figs. 2(d) and 3(d) (closed red squares), and the corresponding enthalpy component of Δ*G*, Δ*H*, in Figs. 2(d) and 3(d) (open brown circles). Notice that the enthalpy, which is derived from Δ*G*+*T*Δ*S cannot* be determined independently. And due to the inherent numerical difficulty in determining free energy differences, the data in Fig 2. 2(d) and 3(d) must be considered suggestive rather than quantitative. In particular, in the region to the left of the first maximum in Δ*G* (vertical dashed line) for each pair going toward the global minimum in Δ*G* (vertical solid line), the data in Figs. 2(d) and 3(d) suggest there is a rise in the entropy component −*T*Δ*S* as a function of decreasing interbase separations. We believe this is likely due to the numerical errors coming from subtracting two rapidly varying Δ*G* at different temperatures from one another. But qualitatively, the data in Figs. 2(d) and 3(d) suggest a local minimum exists in the −*T*Δ*S* term of the free energy at the first maximum of Δ*G* (vertical dashed line), and a local maximum in −*T*Δ*S* exists at the second minimum (vertical dotted line) of Δ*G*, though the precise amplitudes of these are uncertain. It is important to point out that the solvent is the *only* source of entropic effects when considering the base pairing free energy in the stretch direction. This is because the bases are constrained to be coplanar when the free energy profiles are computed, and as a result they have no additional degrees of freedom between them that could give rise to any entropy. Consequently, when the free energy is calculated along the stretch direction in Figs. 1(a), 2(a) and 3(a), *the solvent must be solely responsible for all the observed entropic effects* in Δ*G*(*x*), which must also be entirely contained in the solvent’s contribution Δ*G*_S_(*x*). Therefore, the entropy term −*T*Δ*S* revealed by the data in Figs. 2(d) and 3(d) provide another perspective on the solvent’s role in base pair complementarity, but given the intrinsic uncertainties in these data, we will refrain from overinterpreting them.

## 4. Discussion

### Presence or absence of water impacts different base pairs differently

The computational results presented above paint a complex picture for how the aqueous solvent is actively involved in the pairing interaction between DNA bases. For both Watson-Crick and noncanonical pairs, their free energies do not seem to scale with the expected number of hydrogen bonds. Instead, G|C pairs are only marginally more stable compared to A|T pairs. The aqueous solvent, for every base pair we have examined, exerts an opposing thermodynamic force to counteract the direct hydrogen bonding interactions between the bases. The strength of the solvent’s thermodynamic resistance is substantially stronger in G|C compared to A|T, suggesting the existence of a solvent-induced compensation effect, rendering the overall stability of G|C comparable to A|T. The granular structure of the liquid solvent is imprinted on the base pairing free energy profiles, and nonmonotonic oscillations in the free energy correlate with the computed entropy change in the solvent, which cycles between local maxima and minima as a function of the interbase separation. Stacking and pairing thermodynamic forces, both heavily renormalized by the presence of the aqueous solvent, appear to act approximately additively in their overall effects on the stability of stacked base pairs in duplex DNAs.

Most significantly, our results show that the A|T pair (−6.8 kcal/mol) and the G|C pair (−8.0 kcal/mol) have similar free energies. However, the solvent’s contribution to these free energies is very different for A|T (+4 kcal/mol) compared to G|C (+13 kcal/mol). This solvent resistance acts against the much stronger direct hydrogen-bonding interactions between G|C than A|T, producing a G|C free energy that is ~1.2 kcal/mol more stable than A|T, which as we have shown above, are in line with experimental thermodynamic measurements.

The marginal difference between the thermodynamic stability of a G|C pair compared to A|T invites the obvious question: Is DNA replication governed by the same thermodynamics, and how is fidelity enforced with such a minute free energy difference between the two different Watson-Crick pairs? The answer to the second part of the question is clear from studies of DNA replication – while the melting free energy difference between A|T and G|C pairs is minute, fidelity is enforced because DNA polymerase is able to match A to T and G to C with an accuracy rate of a million to one^49^. The results here show that water is the key. More precisely, the presence or absence of water molecules in the line of contact between the bases is the key that determines matches versus mismatches. For the A|T pair, the solvent’s resistance is worth only +4 kcal/mol, whereas for the G|C pair the solvent’s resistance is much more significant at +13 kcal/mol. For a G|T pair on top of a G|C, the solvent offers little to no resistance at all. Under water, the solvent’s resistance has therefore been delicately tuned against the stronger direct interactions between solvent-exposed G|C versus A|T. However, inside the catalytic core of polymerase where some of these waters are absent, we expect the solvent’s resistance would have been detuned. This detuning inside the dry or drier interior of polymerase produces different selectivity on the G|C pair compared to A|T. Furthermore, any perturbation to this aqueous environment also affects the G|C and A|T pair disparately. In particular, shape matching between DNA bases may take on increased importance in the interior of polymerase, since different approach geometries can exclude different number of waters between the two bases and therefore produce differential effects on the detuning of the solvent’s resistance for matched and mismatch pairs. Because of the significant difference between DNA base pairs in water versus inside polymerase, the thermodynamic operating conditions during DNA replication are likely different compared to naked DNA base pairs in water. We will show below that even a few missing water molecules can produce significant renormalization in the solvent’s resistance.

### Waters recruited to line the gap between bases undergo “freezing”

To visualize how the solvent molecules assert themselves in controlling the pairing interactions between the bases, we have examined the equilibrium distributions of water molecules in the microscopic neighborhood of the bases. Examples are shown in Fig. 4 for the A|T singlet in panel (a) and the GG|CC doublet in panel (b) with their interbase separations corresponding to the first maximum in their respective free energy curves. Using large ensembles of uncorrelated snapshots from our MC simulations, we have computed the average density of water molecules in the vicinity of the bases, on a grid of 0.5Å×0.5Å×0.5Å cells tiling the simulation box. Fig. 4(a) shows densities along the plane passing through the A and T bases on the horizontal plane in green, and densities along the plane perpendicular to the bases on the vertical plane in red, as a multiple of ambient water density. For the A|T pair in Fig. 4(a), densities on the horizontal plane reveal a solvent cavity coinciding with the volume occupied by the bases. Noticeably, there are no water molecules between the two bases. The interbase separation, *x* ~ 4.3 Å, at the first maximum is too narrow for waters to fit. The vertical plane shows the solvent density in red along a plane perpendicular to the bases, cutting through mid-gap between the bases. Clearly, no water molecules are found in the space between the two bases. Note that there is a single moderately high-density spot on the vertical plane, approximately three times the density of ambient water, located above one of the hydrogen bonds in the A|T pair. Other than this, the solvent density appears fairly homogenous in the surroundings of the A|T pair, at the interbase separation *x* ~ 4.3 Å, at the first maximum of the free energy profile Δ*G*(*x*).

**FIGURE 4.**
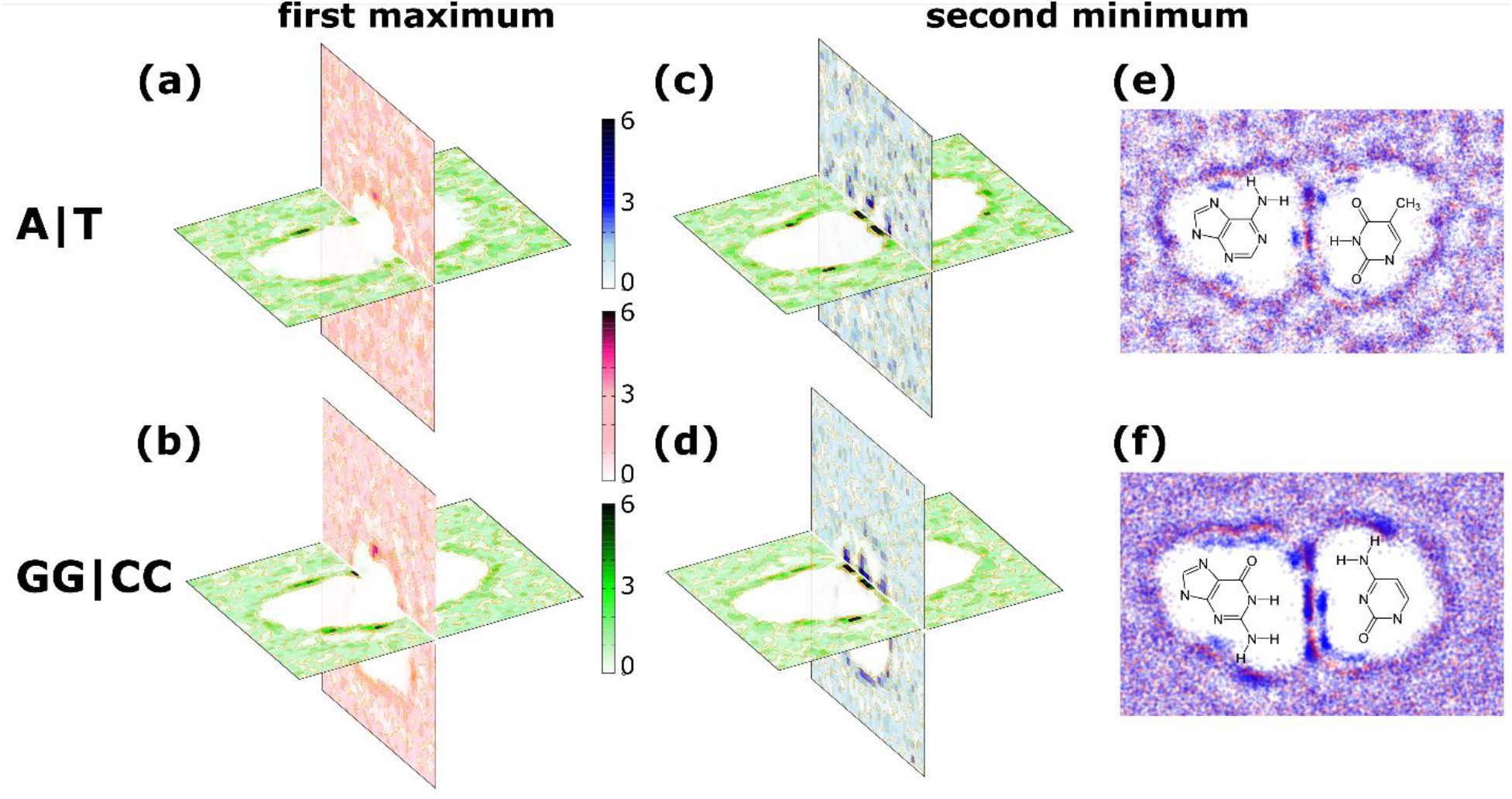
(a) False-color plots showing solvent densities in the vicinity of the A|T base pair along the plane containing the bases (horizontal, green) and the plane perpendicular to the bases (vertical, red), when the A|T separation is at the first maximum of its free energy profile. Color intensities reflect water densities as a multiple of the mean liquid density in the ambient solvent. (b) Solvent densities for the GG|CC pair, at the first maximum of its free energy profile. (c), (d) Solvent densities for the A|T and GG|CC pairs, at interbase separations equal to the second minima in their respective free energy profiles. (e), (f) Scatter plots showing positions of water molecules, with red dots corresponding to O atoms and blue dots to H atoms, from a number of MC snapshots, projected onto the plane containing the bases, relative to the positions of the base atoms. Water molecules appear to orient themselves along the gap between the two bases to try to bridge the hydrogen bonding donors on one base with their corresponding acceptors on the other.

Fig. 4(b) shows densities along the two orthogonal planes for the GG|CC doublet pair. The results are largely similar to the A|T pair, except densities on the vertical plane reveal another cavity below the top-layer G|C bases. This cavity coincides with the volume occupied by the second G|C pair in the GG|CC doublet. Similar to the A|T pair, there is a moderately high-density spot above the bases, adjacent to one of the hydrogen bonds between G and C. There are also several moderately high-density spots along the vertical plane. These are due to strong partial charges situated on the non-Watson-Crick edge of guanine, which attract water, but the densities in these areas do not change significantly with interbase separations.

More distinct features in the solvent density maps show up when the interbase separation becomes large enough to accommodate water molecules between the bases. This is shown in Figs. 4(c) for A|T and 4(d) for GG|CC. For the density shown in Fig. 4(c) for the A|T pair, the interbase separation is x ~ 5.5 Å, which corresponds to the second minimum in its Δ*G*(*x*) curve. Along the horizontal plane, two very high-density spots are observed between A and T, where no water was found between the two bases previously when the interbase separation was at the first maximum. The positions of these high-density regions coincide with the two hydrogen bonds found in the A|T base pair. Corresponding high-density spots are also observed on the vertical plane in the A|T pair. These indicate that two water molecules are attracted to the gap between the two bases, which provides a very tight volume into water molecules are recruited. The densities inside these regions of tightly sequestered water are roughly 10-fold over ambient water density. In order to produce this kind of density enhancement, the volumes available to the waters that are recruited into these confined regions must be ~ 10-fold smaller than ambient water. From this, we estimate that the entropic penalty associated with each of these confined waters must be approximately *R*ln(10), where *R* is the gas constant. At 37°C, this corresponds to a free energy cost of ~ 1.4 kcal/mol per confined water trapped between the two bases.

The picture for the GG|CC doublet in Fig. 4(d) is similar. But instead of two high-density spots between A and T in Fig. 4(c), there are three high-density spots between G and C. These correspond to three water molecules trapped between hydrogen bond donors and acceptors between G and C. The density enhancement in these high-density regions over ambient water is similar to what was found in the A|T pair, leading to an entropic penalty of ~ 1.4 kcal/mol of free energy per confined water molecule. To confirm that these high-density regions are indeed due to water molecules trapped in the space between the two bases, Figs. 4(e) and (f) show scatter plots of the oxygen (red dots) and hydrogen (blue dots) atoms belonging to water molecules located in the vicinity of the bases, for A|T and GG|CC, respectively, from a number of uncorrelated solvent configurations collected over the simulations. Clearly, the high-density regions between the bases correspond to distinct water molecules. Each of these trapped waters appears to be using one of its O-H bonds to try to bridge the gap between a hydrogen bond donor on one base and its corresponding acceptor on the other base. The solvent, in order to accommodate this, undergoes a structural reorganization to recruit water molecules to line the gap between the bases, in an apparent attempt to try to minimize the solvent’s enthalpy, simultaneously producing a confinement for these water molecules, resulting in suppression of the solvent’s entropy. Within this interpretation of the data, the thermodynamic driving force behind solvent reorganization is the tendency of the solvent to minimize interaction energy between the bases, while the compensating factor is the solvent’s decreasing entropy, similar to freezing. Based on enthalpy entropy compensation in DNA doublet melting free energies, Petruska and Goodman^15,16^ have also postulated that water molecules within the solvation shell can contribute to the stability of each double in a way that may be related to the freezing of these waters.

When the interbase separation between the two bases decreases beyond the second minimum, the gap between them is no longer able to accommodate water. As a result, water molecules must be evacuated from the gap between the bases, releasing them from a state of strong confinement back into a higher-entropy state within the ambient solvent background. Because of this, the solvent’s entropy is expected to increase. This is consistent with the decrease observed in the entropic component −*T*Δ*S*(*x*) in Figs. 2(d) and 3(d), to the left of the second minimum

### A handful of near-field waters account for the majority of the solvent’s resistance

While we have argued that the presence or absence of waters in the interior of DNA polymerase may be used as a selectivity filter to favor matched versus mismatch base pairs, the obvious question is how many water molecules actually make up the solvent’s resistance and how many of these must be excluded from the solvent for this selectivity effect to be enforced. In the last section, we have observed that distinct water molecules can freeze into the interface between the two bases at very specific interbase separations, but we have not been able to identify other clearly distinguishable waters solely from the solvent’s density in other regions of interbase separations. Therefore, to try to identify those water molecules that are responsible for the solvent’s resistance, we carried out the following experiment. For each configuration along the MC trajectory, we searched for water molecules around each base and deleted waters that were within 3.0 Å from any Watson-Crick donor or acceptor atom, and with these waters removed re-evaluated the gradient of the total energy between the bases. Gradients collected from MC configurations without these “near-field” waters (-water) were then used to calculate the ensemble-averaged thermodynamic force between the two bases. These were compared against the forces calculated with all waters in place (+water), and using the difference between −water and +water, we were able to determine the net contribution due to the near-field waters. The integral of this then produced the free energy of the solvent due to these near-field waters alone, and the results are shown in Fig. 5.

**FIGURE 5.**
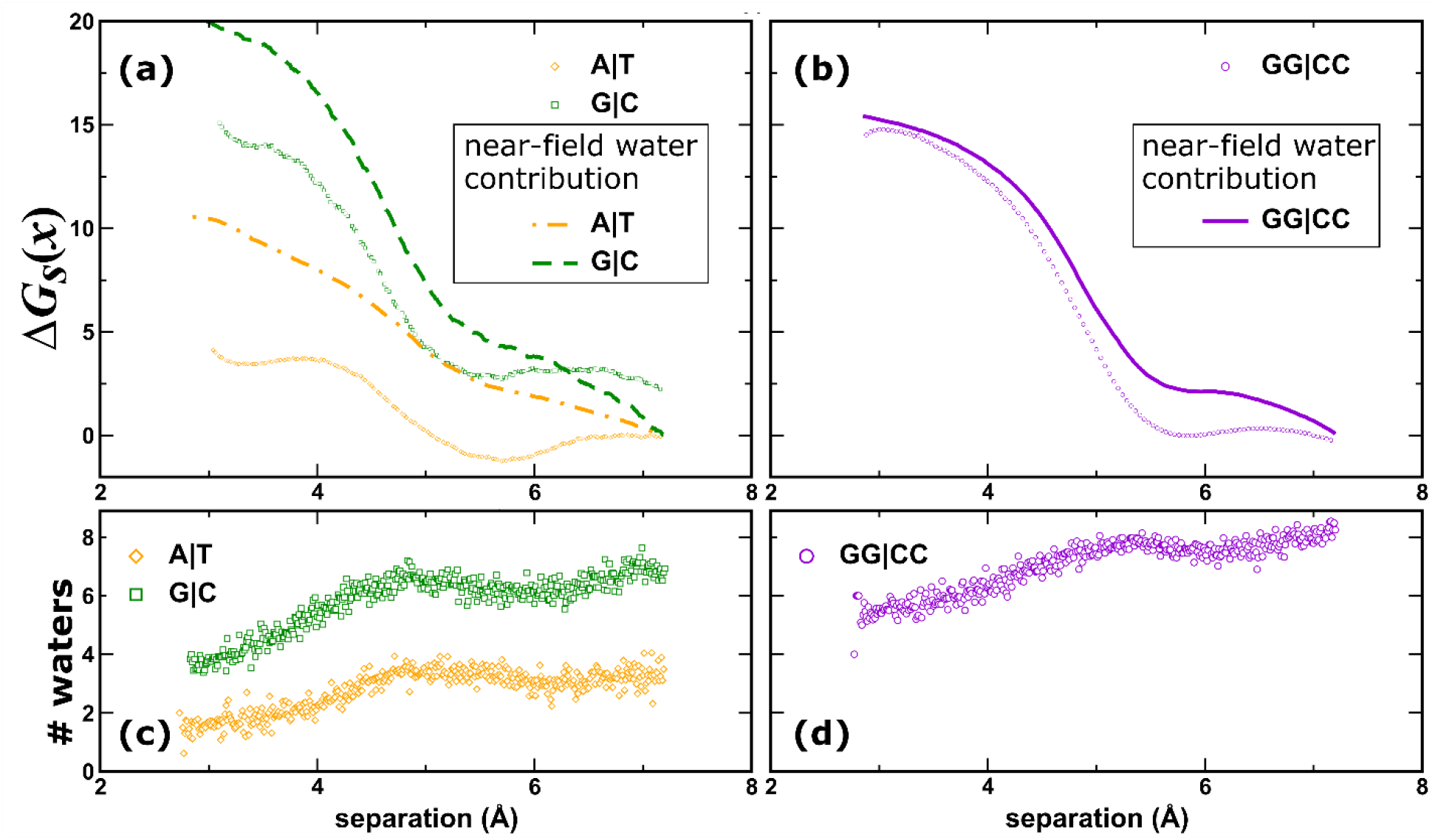
(a) The contribution of near-field waters to the solvent’s resistance to base pairing: A|T (dotted dashed line) and G|C (dotted green line), compared to full solvent results from Fig. 2(b): A|T (open orange diamonds) and G|C (open green squares). (b) The contribution of near-field waters to the solvent free energy for GG|CC (solid purple line) compared to full solvent results from Fig. 3(b) (open purple circles). (c), (d) Average number of near-field waters in each A|T (open orange diamonds), G|C (open green squares) and GG|CC (open purple circles) as a function of base separation, showing that a small number of near-field waters account for the majority of the solvent’s resistance to base pairing.

Fig. 5(a) shows the near-field waters’ contribution to the solvent free energy for A|T as the orange dotted dashed line, and for G|C as the green dashed line. The free energies in full solvent from Fig. 2(a) are shown in Fig. 5(a) for comparison (AT +water: open orange diamonds, GC +water: open green squares). Fig. 5(b) shows the same results for GG|CC, with the near-field waters’ contribution as the purple line, and the full solvent as the open purple circles. For the singlet A|T and G|C pairs, Fig. 5(a) shows that the near-field waters collectively produce a stronger destabilizing force against base pairing compared to the full solvent, which implies that the far-field waters must provide a mildly stabilizing force for the base pair, ~ −6 kcal/mol for both A|T and G|C. For the GG|CC doublet, Fig. 5(b) shows that the near-field waters’ contribution (solid purple line) is almost equal to the full solvent (open purple circles) in its entirety. Figs. 5(c) and (d) show the average number of these near-field waters for A|T (open orange diamonds), G|C (open green squares) and GG|CC (open purple circles) as a function of interbase separation. The number of these near-field waters are generally quite small. In the case of the stacked GG|CC doublet pair, this small set of near-field waters turns out to make up the majority of the solvent’s resistance to base pairing. Since the formation of the GG|CC doublet most closely reflect the expected approach geometry when the copy strand is synthesized within the catalytic core of polymerase, we believe that near-field waters largely control the aqueous solvent’s resistance to base pairing, and the exclusion of one of more of these near-field waters may be used in the interior of polymerase to select for matched base pairs against mismatches.

## 5. Conclusion

We have conducted a series of large-scale Monte Carlo simulations to study base pairing in explicit water to examine the solvent’s role in determine the strength of DNA base pairs. We quantified the free energy of singlet G|C, A|T and G|T pairs, as well as a doublet GG|CC pair. For singlet pairs, the free energy of G|C is only marginally more stable than A|T pair or G|T. We have explicitly computed the solvent’s contribution to the base pairing free energy in each and found that the aqueous solvent exerts an almost exclusively destabilizing thermodynamic force to counteract the direct hydrogen bonding interactions between the bases. The strength of the solvent’s thermodynamic resistance is substantially stronger in G|C compared to A|T, suggesting the existence of a solvent-induced compensation effect, rendering the overall stability of G|C very similar to A|T under water. The granular structure of the liquid solvent is imprinted on the base pairing free energy profiles, and nonmonotonic oscillations in the free energy correlate with the computed entropy change in the solvent, which cycles between local maxima and minima as a function of the interbase separation. Stacking and pairing thermodynamic forces, both heavily renormalized by the presence of the aqueous solvent, appear to act approximately additively in their overall effects on the stability of stacked base pairs in duplex DNAs. Using equilibrium density profiles of the water in the vicinity of the bases, we correlated the decrease in solvent entropy around the second minimum in the pairing free energy profile with a “freezing” phenomenon where water molecules from the solvent are recruited to line the interface between the two bases in the pair, in order to minimize the solvent’s enthalpy. As the interbase separation decreases, these frozen water molecules are expelled from the interface when they can no longer fit within the gap between the bases, producing a “thawing” behavior where entropy increases again. As the solvent undergoes reorganization during these freeze-thaw events, the entropy of the solvent cycles.

These results demonstrate that while the G|C pair is only marginally more stable than A|T in water, we surmise the selectivity filter employed by polymerase to enforce replication fidelity may be controlled by the presence or absence of water molecules inside its catalytic core. The solvent’s thermodynamic resistance to pairing is almost exclusively destabilizing, but the effect is disproportionately larger for G|C than A|T, and this might be used to select for matched base pairs against mismatches. A handful of near-field water molecules associated with the Watson-Crick donor/acceptor atoms on the bases account for the majority of the selectivity effect. Since the approach geometry during the pairing of the bases can affect the number of these near-field waters, shape matching between DNA bases can also affect replication fidelity.

## Acknowledgements

This material is based in part upon work supported by the National Science Foundation under Grant Numbers CHE-0713981 and CHE-1664801. The authors thank Myron F. Goodman for helpful discussions and comments.

